# High-throughput phenotyping with deep learning gives insight into the genetic architecture of flowering time in wheat

**DOI:** 10.1101/527911

**Authors:** Xu Wang, Hong Xuan, Byron Evers, Sandesh Shrestha, Robert Pless, Jesse Poland

## Abstract

**Background:** Precise measurement of plant traits with precision and speed on large populations has emerged as a critical bottleneck in connecting genotype to phenotype in genetics and breeding. This bottleneck limits advancements in understanding plant genomes and the development of improved, high-yielding crop varieties.

**Results:** Here we demonstrate the application of deep learning on proximal imaging from a mobile field vehicle to directly score plant morphology and developmental stages in wheat under field conditions. We developed and trained a convolutional neural network with image datasets labeled from expert visual scores and used this ‘breeder-trained’ network to directly score wheat morphology and developmental stages. For both morphological (awned) and phenological (flowering time) traits, we demonstrate high heritability and extremely high accuracy against the ‘ground-truth’ values from visual scoring. Using the traits scored by the network, we tested genotype-to-phenotype association using the deep learning phenotypes and uncovered novel epistatic interactions for flowering time. Enabled by the time-series high-throughput phenotyping, we describe a new phenotype as the rate of flowering and show heritable genetic control.

**Conclusions:** We demonstrated a field-based high-throughput phenotyping approach using deep learning that can directly score morphological and developmental phenotypes in genetic populations. Most powerfully, the deep learning approach presented here gives a conceptual advancement in high-throughput plant phenotyping as it can potentially score any trait in any plant species through leveraging expert knowledge from breeders, geneticist, pathologists and physiologists.

## BACKGROUND

Limitations in phenotyping are widely recognized as a critical constraint in genetic studies and in plant breeding [1, 2]. Initial developments in high-throughput phenotyping (HTP) have focused on direct sensor or image measurements to extract proxies for traits of interest such as vegetation indexes from spectral reflectance [3, 4] or plant height from digital elevation models [5]. While lending great insight to plant processes, this first-generation of HTP is limited in assessment of ‘complex’ traits such as plant morphology or growth stage that that cannot be assessed by a linear function of pixel values. While these complex morphological and developmental features are readily distinguished by a trained eye, the assessment of these phenotypes with high-throughput platforms is challenging.

Deep learning has emerged as a powerful machine learning approach that takes advantage of both the extraordinary computing power and very large datasets that are often now available [6]. Deep learning bypasses the need to explicitly define which features are most useful or needed for data analysis. Instead deep learning discovers a complete end-to-end process to map data samples to outputs that are consistent with the large, labelled datasets used for training the network. For image analysis tasks, convolutional neural networks (CNNs) learn this end-to-end mapping by optimizing for many layers of filters. The first filters are easily interpreted as image features (e.g. detecting edges, bright points or color variations), and subsequent layers are increasingly complicated combinations of earlier features. When there is sufficient training data, CNNs dramatically outperform all alternative existing methods for image analysis. For benchmark classification tasks attempting to label which of 1,000 different objects are in an image, results have increased from 84.6% in 2012 to 96.4% in 2015 [7]. Based on this impressive performance of the latest CNNs, we hypothesized that this deep learning approach could be an equally powerful tool for scoring subtle, complex phenotypic differences directly from images in segregating plant populations.

## DATA DESCRIPTION

### Field-based high-throughput imaging of wheat plots

To advance high-throughput phenotyping of complex morphological and developmental traits, we developed a high-clearance field vehicle [8] equipped with an array of DSLR cameras collecting geo-positioned images (**Figures 1a & 1b**). This platform was deployed across wheat field trials in 2016 and 2017. Each year we grew two trials, one recombinant inbred line (RIL) population from a cross between wheat cultivars ‘Lakin’ and ‘Fuller’ and a panel of diverse historical and modern winter wheat varieties consisting of a total of 1398 plots each year. We captured over 400,000 proximal images of the wheat canopies throughout the growing seasons in 2016 and 2017. These images were geo-referenced and 135,771 and 139,752 of the images were assigned to individual field plots in 2016 and 2017, respectively, based on surveyed coordinates of the field plots and geo-tagged images (**Fig. 1, Supplementary Fig. S1**). This approach enabled high-throughput proximal imaging on an individual plot level (1.5m × 2.4m plot size). Concordant with imaging, field plots were visually scored for percent heading and spike morphology of awned or awnless. To generate a large collection of labeled images suitable for deep learning, the images were labeled with visual ‘breeder scores’ taken on the same respective plots at the same time point (**Fig. 1c**). The data set(s) supporting the results of this article are available in the *Giga*DB repository, [persistent identifier].

## ANALYSIS

### Development of convolutional neural networks

To assess plant features that cannot be measured directly by sensors with the high-throughput platform, we developed a CNN network that could be trained using these geo-positioned images labeled with visual scores and subsequently automatically score plots for the phenotypes of interest. As a starting point, we first approached the qualitative trait of awn morphology (**Supplementary Fig. S2**).

An initial challenge in the development of the CNN was memory constraints that limit the networks to analyzing relatively small images, but the images were captured at very high resolution. Because the relevant image features are quite small (e.g. wheat awns at 1-2mm width) reducing the size of the image would make these features invisible. We therefore, cropped the images into a three by three grid of patches at 224 × 224 pixel size. To build the full ‘WheatNet’, we then extended the CNN architecture that analyzes images with a small additional network that combined features from the nine patches to create a consensus estimate for the image (**Fig. 1d**).

We used this developed CNN architecture in a training-validation-testing approach to predict the awn phenotype in the diverse panel of inbred lines in which there were awned and awnless variants. The training and validation images were from this diversity panel evaluated in 2017 with 700 plots, of which 29 plots were awnless and 671 were awned. Model training used 2000 images for awned plots and 1800 images for awnless plots. As a validation dataset, we sampled 70 plots from the awned and 5 plots from the awnless and left the remaining plots as the training data. We validated the WheatNet on a set of 300 images each from awned and awnless plots. On this set, the network prediction matched the breeder assessment at 99.20% on the training set and 98.61% on the validation set.

To test the WheatNet for scoring awn morphology, we applied the network trained with data from 2017 to test images from field trials of the diversity panel in 2016 which contained 12,504 images from 675 awned plots and 32 awnless plots. On an individual image prediction accuracy was 98.9% for awned and 98.7% for awnless phenotypes (**Supplemental Table S1**). As many images were captured for each plot, we applied a plot-level consensus voting which increased the accuracy to 99.7% for awned and 100% for awnless. Strikingly, we observed that only two plots were inconsistent between visual scoring and the CNN and that these two plots were the same variety (‘MFA-2018’) across both field replications. Further inspection showed that this variety was heterogeneous and has an ‘atypical awnlette’ phenotype, indicating that the CNN was able to detect subtle atypical phenotypes that were lost or ignored in the human scoring.

### Measurement of percentage heading

Flowering time is a critical trait under intense selection in natural and selected breeding populations. Due to tightly closed flowers in wheat, spike emergence (heading time or heading date) is used as a close proxy for breeding and genetics. Observing initial proof of concept for using deep learning to score a simple Mendelian morphological trait, we extended this approach to a more complex problem of developmental phenotypes using time-series imaging. To score the heading date of wheat, which is classically defined as the date in which spikes (ears) have emerged from 50% of the tillers[9], we applied the CNN to scoring percentage heading over longitudinal image datasets. For the initial trait scoring of percent heading we used the same CNN training approach as was used for predicting the wheat awn phenotype and trained the network with two years of image data from the diversity panel. An additional feature of scoring percentage heading makes this problem different from standard classification problems, in that the breeder labels are given in 10% increments, and there may be some inconsistency in these labels. To address this, we modified the algorithm that trains the CNN to give partial credit for predictions that are either 10% and 20% different from the correct label.

On the classification problem for percent heading the network prediction exactly matched the ten percentile classifications from visual assessment on 45.12% (training set) and 41.27% (validation set) of the images, which is much better than a random guess at 9.10% (1 out of 11 classes). The confusion matrix for training, validation and testing the CNN for predicting percent heading showed clear diagonal patterns indicting linear consistency between the observed and predicted values (**Supplemental Figure S3**). Though there was lower accuracy in the testing phase, the diagonal linear pattern remained consistent. From these observations, we had strong evidence that the CNN is accurately scoring percentage heading for images across the range of heading values and throughout the season. Following this conclusion, we applied the CNN trained on the diversity panel to score percentage heading and calculate heading date for a biparental RIL population and determine if genotype-to-phenotype associations could be detected from phenotypes scored directly by deep learning.

To translate the time-series imaging and CNN phenotypic measurements into a single time point for assessment of heading date, we applied a logistic regression to the percent heading measurements for each individual plot (**Fig. 2a**). From the logistic regression fit, we then found the date intersecting the regression at 50% and called this time point as the heading date commensurate with the classical definition of 50% of heads emerged from the boot. Applying logistic regression individually to each plot we obtained heading dates from the CNN that were highly accurate with the heading date assessed directly from visual scoring (**Fig. 2b**). We observed over 57% and 88% of heading date measurement were within one day and two days, respectively, with a mean absolute error of 0.99 and mean root square error of 1.25 days. Slope of the regression between the visual and CNN measurements of 1.02 indicated lack of bias from the CNN. Reflecting accurate phenotypes under strong genetic control, the broad-sense heritability for heading date was very high when measured by both visual (*H*^*2*^=0.982) and CNN (*H*^*2*^=0.987) scoring.

An interesting novel phenotype that can be assessed with this time-series approach is the developmental rate of flowering (heading) progression within an inbred line. This rate of heading measure can be derived from the slope of the logistic regression. Measuring the slope for each inbred line, we found heritable genetic variance for the rate of heading (*H*^*2*^_*VISUAL*_=0.621, *H*^*2*^_*CNN*_=0.514), indicating that this developmental rate phenotype is also under genetic control. As the rate of heading might simply be an artifact from the heading date *per se*, we tested the correlation between timing and rate of heading and found weak negative correlation (*r* = −0.19, *p-value* < 0.001). Looking at RILs within the normal early range of heading date (e.g. before May 5^th^; Day 125) we found no significant correlation (*r* = 0.078, *p-value* = 0.079) suggesting that the rate of heading is indeed under independent genetic control from heading date.

### Genetic analysis of flowering time

Following the measurement of heading date using the neural networks, we sought to determine the utility of phenotypes scored directly from deep learning to uncover the genetic basis of the variation in flowering time present in the biparental population. Though the parental lines ‘Lakin’ and ‘Fuller’ are very similar in heading, the progeny showed vast transgressive segregation and also segregation distortion indicating some underlying epistatic gene action (**Supplementary Fig. S4**). We implemented a genome-wide scan of 8237 genotyping-by-sequencing markers and found strong associations for *PpD-D1* and *PpD-B1* as well as associations on 5B and a novel QTL positioned on at the distal end of chromosome 1B (**Fig. 2c**). Suspecting epistatic gene action based on the phenotypic distribution, we tested all significant markers for putative epistatic interactions and found strong interactions between *PpD-D1* and *Ppd-B1* as well as between *PpD-D1* and the locus on 1B (**Fig. 2c**). Interestingly, the modeling of the interactions between all three loci removed the main effect of the 1B locus *per se*, with this locus having an opposite effect in the presence of *PpD-D1* early (insensitive) allele (**Fig. 2d**).

While we found heritable genetic variance for the rate of heading, we were not able to find any genetic association within this population (**Supplementary Fig. S5**). Implicating missing heritability for the rate of flowering suggested a diffuse genetic architecture of many small effect alleles. We therefore tested whole-genome polygenic models (BayesA and G-BLUP) to capture the genetic variance for the rate of heading. We ran 100 replications of cross-validation predicting 10% masked phenotypes and were able to model 18 to 25% of the heritable genetic variance.

## DISCUSSION

From this study, we have demonstrated that deep learning with breeder-trained neural networks from proximal field-based imaging can accurately score plant morphology. When applied to time-series image datasets this approach can likewise accurately score developmental stages such as flowering time. Furthermore, these machine vision phenotypes can be used directly to uncover genetic determinants in the populations, connecting genotype to phenotype in the same way as classical approaches to phenotyping.

A significant advancement of the approach presented here is that there is no additional time investment in developing the labeled image set for training. Many applications applying deep learning for image based phenotyping require extensive annotation of the training image sets, such as annotating plant features of interest prior to network training [10, 11]. As demonstrated in this study, HTP image datasets from field trials can be labeled through direct imputation of visual scores routinely collect in the field by breeders. This approach can develop a labeled image training dataset for any phenotype of interest without any further human input.

Though the images are labeled, and hence the network subsequently trained, by experienced individuals, there are inherent limitations and bias associated with visual scoring of any type [12]. Just as expert breeders, pathologists and physiologists can disagree among themselves on how to classify subtle phenotypic differences, the CNN developed here notably had a case of consistent disagreement with the visual scores from the expert that was actually used to train it. Indeed, it has been shown in different fields how deep learning can surpass the accuracy of experts [13]. Moving beyond the input of a single person, deep learning for high throughput phenotyping has the potential power to synthesize the consensus knowledge of an entire community of experts through training on shared datasets. When combined with high-resolution imaging that is becoming more easily acquired from unmanned aerial systems, the collection of this level of image data across many research groups and breeding programs could develop robust training sets for all phenotypes of interest along with the built-in feature of consensus from many experts.

## POTENTIAL IMPLICATIONS

The first generation of high-throughput plant phenotyping has focused on sensor and image features that can be directly mapped to plant phenotypes, but remains limited on exploring the full scope of phenotypic variants. Conceptually, this deep learning approach for the next-generation of high-throughput phenotyping can be extended to any trait of interest in any species for which high-resolution imaging and expert scoring of the phenotypes can be obtained. This development in high-throughput plant phenotyping can enable breeders and geneticists to measure complex phenotypes on the size of populations needed to understand gene function on a genome-wide scale and uncover genetic variants to develop vastly improved varieties for a future of food security.

**Figure 1.**
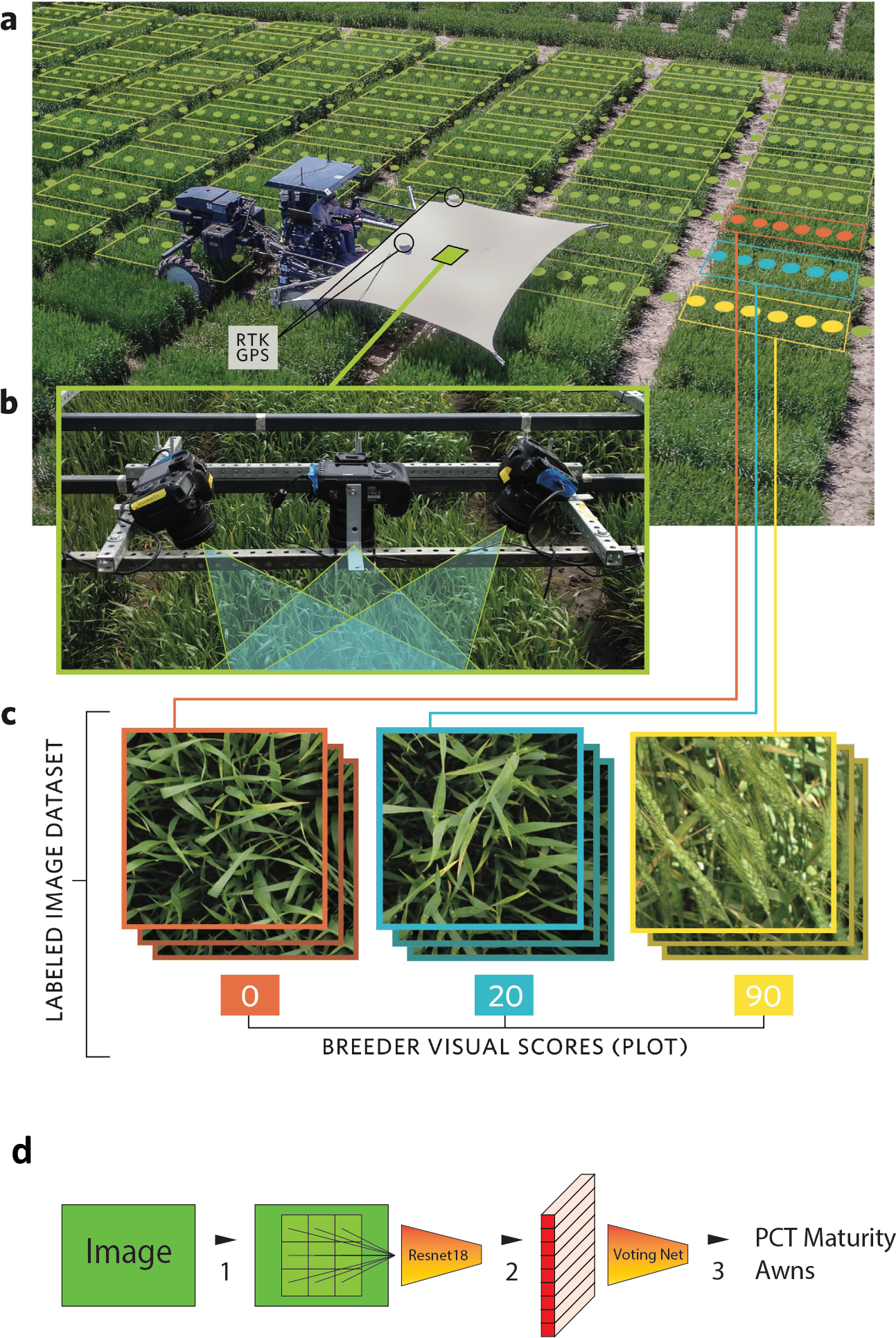
Phenotyping platform and “breeder-trained” image data sets in this study. (**a**) Aerial view of field-based high-throughput phenotyping platform deployed in current study traversing wheat plots with superimposed example representation of imaging positions and example field plot boundaries. (**b**) imaging array of multiple DSLR cameras deployed on the phenotyping platform to collect geo-referenced proximal images of the wheat canopy. (**c**) combination of images assigned to respective field plots and merged with visual breeder scored to develop the labeled imaged dataset for training the neural networks. (**d**) Schematic of the “WheatNet” neural network developed for classifying cropped image patches followed by census voting network for the whole image.

**Figure 2.**
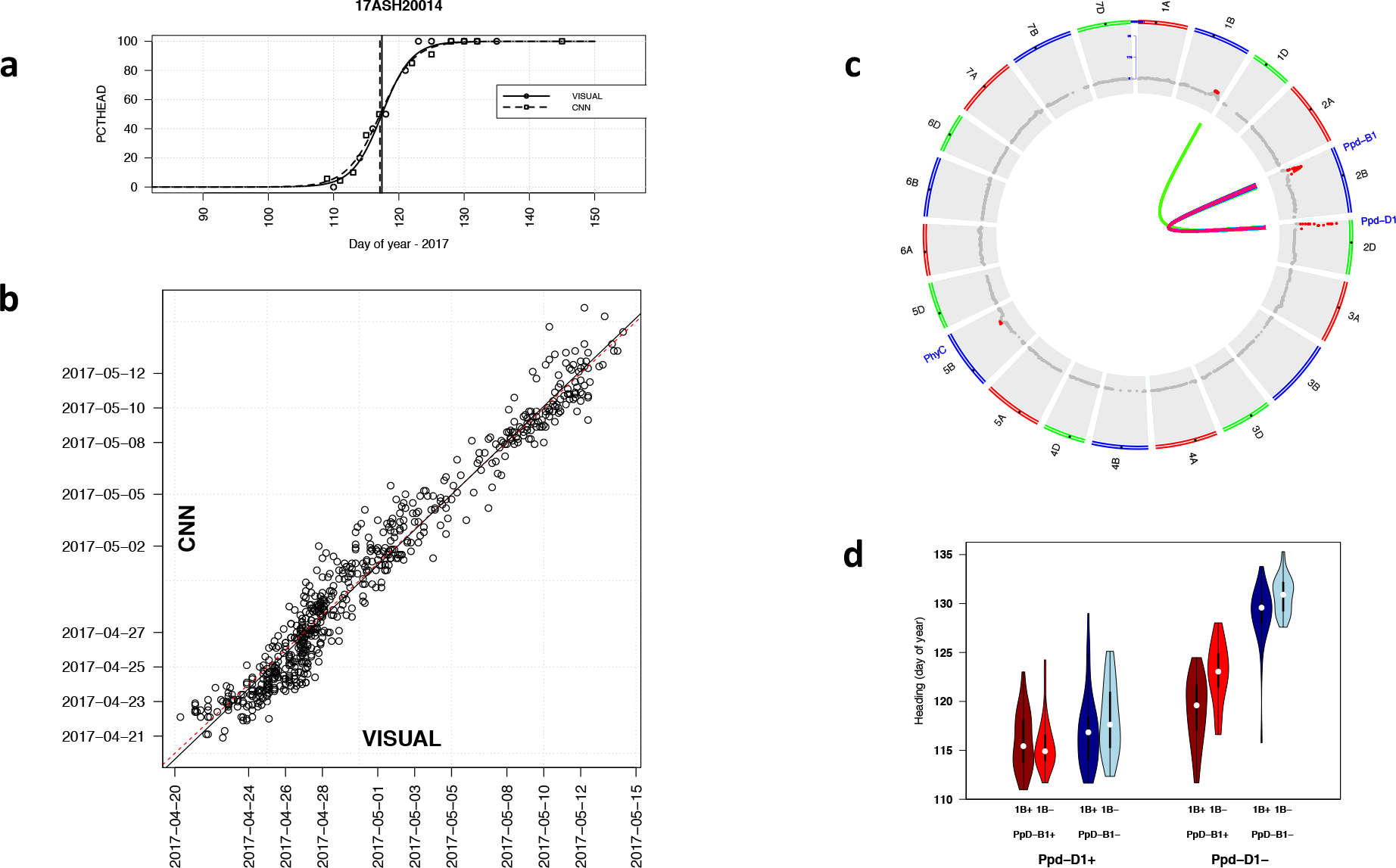
Genetic analysis of heading date scored by deep learning. (**a**) Example of logistic regression to score heading date from time series measurements for a single field plot (17ASH20014). Shown are visual scores (circles) and predictions from the convolutional neural network (squares) and the respective fitted regression lines. The 50% interaction of the regression was identified as scored as the heading date. (**b**) Heading date compared between visual scoring and convolutional neural network (CNN) from logistic regression applied individually to 676 field plots for percentage heading from time-series observations (visual) and time-series imaging (CNN). Scoring (VISUAL) and imaging dates (CNN) are shown on the axes. (**c**) Genome wide association testing for heading date measured by deep-learning on the Lakin x Fuller recombinant inbred line testing for 21 wheat chromosomes. Markers above Bonferroni multiple-test correction threshold shown in red. Epistatic interactions (internal connections) for two-gene interaction model. Significant interactions at Bonferroni correction show with heat-colors indicating significance level (LOD range from 4.97 (blue) to 22.4 (red)). (**d**) Phenotypic distributions of genotypes at loci showing significant epistatic interactions including Ppd-D1, Ppd-B1 and QTL identified on Chr. 1B in the Lakin x Fuller RIL population. Plus (+) and minus (−) indicates genotypes with or without the early allele.

## METHODS

### Development of field-based high-throughput platform for image collection

A phenotyping mobile unit (PheMU) [8] was developed to measure winter wheat phenotypes at Kansas State University (KSU), Manhattan, Kansas. PheMU was constructed on a high-clearance sprayer (Bowman Mudmaster, Bowman Manufacturing Co., Inc., Newport, Arkansas, USA) with a height-adjustable sensor boom to capture images throughout the growing season from consistent distance from the canopy. An imaging array including multiple digital single-lens reflex (DSLR) cameras (EOS 7D, Canon) was carried by PheMU to capture high-resolution crop images. For georeferencing images, two GNSS antennae (AG25, Trimble; Sunnyvale, California, USA) were installed at each end of the sensor boom and connected to an RTK GNSS receiver (BX982, Trimble; Sunnyvale, California, USA). A laptop computer was used to control cameras, collect images, and log positioning data. In addition, to reduce the shadows on the canopy and capture images in a balanced light condition, a rectangle-shape shade sail (Kookaburra OL0131REC, Awnings-USA, Camanche, Iowa, USA) was mounted over the sensor boom.

### Plant Materials and Field Experiments

Two populations were used in this study, 1) a recombinant inbred line (RIL) population consisting of 318 RILs developed from single seed descent to the F5 generation from a cross between U.S. winter wheat varieties ‘Lakin’ and ‘Fuller’, with seed for field trials increased from a single plant in the F5 generation, and 2) a diverse panel of winter wheat inbred lines (Diversity Panel) that has been previously described consisting of 299 winter wheat lines of historical and current cultivars [14]. In the present study, we augmented these lines with an additional 41 newly released cultivars (**Supplementary Data S1**).

Field trials were planted at the Kansas State University Ashland Bottoms research farm (39.131595, −96.619524) on 2015-10-10 and 2016-10-18 for the Lakin x Fuller and 2015-10-20 and 2016-10-19 for the diversity panel for the two years, respectively. Trials were planted in two replications of an augmented incomplete design with one check plot per block of ‘Lakin’ and ‘Fuller’ for the Lakin x Fuller population and ‘Everest’ for the AM Panel (**Supplementary Data S1**)

Phenotypic measurements for awns and percent heading were visually scored and recorded using FieldBook [15]. Awn morphology was scored on the diversity panel according to crop ontology CO_321:0000027 (www.cropontology.org). The Lakin x Fuller population is completely awned. Percent heading was recorded at two to four day intervals through the season scored corresponding to development stages of Zadoks [9] 49 to 59 (Crop Ontology CO_321:0000476). Percent heading was recorded as the percentage of heads emerged from the boot to give a direct indicator with multiple linear classes from 0% to 100% of heading progression to model with the deep learning. A score of 50% heading was given consistent with the standard visual observation of heading date when 50% of the spike is emerged on 50% of all stems (Crop Ontology CO_321:0000840).

### Field Mapping and Image Collection

Cameras in the imaging array were set to capture proximal images of wheat plots in nadir and off-nadir view angles, and from different parts of each plot. The PheMU was operated at 0. to 0.5 m/s with the cameras positioned at 0.5m above the canopy. Each DSLR camera was triggered to take mid-size JPEG images (8 megapixels) in 1.25Hz by a C# program using the Canon EOS Digital SDK (EDSDK v2.14, Canon, JPN). (github.com/xwangksu/CamControl). To capture unblurry and focused images on a mobile platform, the camera was set using manual focus, 1/500-second shutter speed, and F-5 aperture. The camera ISO was adjustable according to the light condition at the beginning of each image acquisition. Each image was directly transferred from the camera to the laptop computer. Image file names and the time stamps when captured were logged in a text file for subsequent georeferencing. Images were then georeferenced and positioned to individual field plots using the approach of Wang *et al*. [16]. The boundary coordinates of each field plot were delineated in Quantum GIS (QGIS, www.qgis.org) using an orthomosaic field map generated from aerial images using the approach of Haghihatilab *et. al.* 2016 [5]. Images inside each plot boundary were geotagged with designated plot IDs to be linked with the genotype information (**Supplementary Dataset**).

### Neural Networks

We followed the standard approach of starting with an existing network pre-trained on the Imagenet dataset [17] and fine-tuning that network to give optimal performance for our task. We subsequently made slight modifications to standard CNN to optimize the network for the phenotyping task. For the baseline model we used Resnet [18], which has been previously used for many applications including people re-identification [19] and flower species identification [20].

#### Training data preparation and size

For the awned phenotype, training and validation images were from 2017 association mapping panel (AM Panel) containing 700 plots, of which 29 plots were awnless and the remaining were awned. As a validation dataset, we reserved 70 plots from the awned and 5 plots from the awnless and sampled images from the remaining plots as the training data. For the training set, we used 2000 images for awned plots and 1800 images for awnless plots. The “WheatNet” network was trained with the following parameters; mini-batch stochastic gradient with a batch size of 44. The learning rate is initialized at 0.01 and was reduced by 80% every 5 training epochs. Training continued for 30 epochs.

For scoring heading percentage the training dataset is from the AM-Panel image set from 2016 and 2017 which consists 711 plots. In these plots, the training set contains 611 randomly sampled plots, and the validation set contains the remaining 100 plots. Each plot was imaged on multiple dates, and on each date, the multiple images were taken of each plot. Each plot was assigned one of 11 classes, corresponding to a visual score of percentage heading of 0, 10, 20, … 100. To create a training dataset, images from each plot were randomly selected to get 2000 images per class. A validation data set was sampled from images from the 100 plots from the diversity panel by randomly selecting 200 images for each class.

Resnet, and most convolutional neural networks, are limited in the size of the image they can analyze. We break the large images captured by the tractor system into 3 by 3 blocks (patches). Each of the 2000 images per class give 9*2000 patches per class. Therefore, overall we train with 198,000 image patches from 22,000 images. The test dataset comes from the Laken-Fuller RIL population in 2017. This dataset contains about 80,000 images per class, and all images came from 676 plots and germplasm that were never seen in the test or validation data.

To develop a more robust CNN for heading percentage we developed two important modifications of the base network.

#### *Modification 1:* Error function that gives some credit for “off by 10-20%” results

The output of a CNN can be viewed as a probability distribution of classes with close percentage classes being more similar. Meanwhile, a given visual labeled maturity percentage can have a +/−10% or +/−20% offset mislabeled condition. In our data, the percentage of the mislabeled images which has offset 10% is about 10% in each class, and the mislabeled images which has offset 20% is about 5% in each class. So, the target label is modeled as a distribution that the labeled class has the value 0.7 for correct class, 0.1 for 10% off and 0.05 for 20% off respectively and the remaining classes have value 0, which ensure the sum of all class probability is 1. The error function calculates the average mismatch of the probability of output and the target distribution, which calculate the absolute difference of each class value between output and the target value.

#### *Modification 2:* WheatNet

In order to keep as much of the full details of the images as possible, which is sensitive to maturity classification, the modified architecture keeps the input image with the resolution of 672 by 672 pixel. The main idea of the design of the architecture is to mimic how experienced individuals assign a phenotype (e.g. the maturity percentage) from a wheat plot or image, by taking a consensus from viewing all part of the images. To capture this feature of visual scoring in the CNN, the network classified the maturity percentage of each image patch and then summarizing all predictions and giving out the total prediction. Full detail of each layer in the Wheat Net is included as **Supplementary Table S3**.

### Genetic Analysis

RILs from the Lakin-Fuller population were genotyped using genotyping-by-sequencing with two-enzymes, *Pst*I and *Msp*I [21]. Two sets of libraries for the RILs and replicated samples of the parents were made in 95-plexing and 190-plexing and sequenced with Illumina HiSeq2000 and NextSeq, respectively. Single nucleotide polymorphisms (SNPs) were called using TASSEL 5 GBS v2 pipeline [22] anchored to the IWGSC wheat genome v1.0 assembly (https://wheat-urgi.versailles.inra.fr/Seq-Repository/Assemblies) with the following parameters: -mnQS 10, enzyme PstI-MspI, -c 20, -minMAPQ 20, and -mnMAF 0.1. Unique sequence tags were mapped to the wheat reference genome (Chinese Spring) using bowtie2 [23] with the following settings: --end-to-end -D 20 -R 3 -N 0 -L 10 -i S,1,0.25. SNPs passing at least one criteria were recovered: inbreeding coefficient of at least 0.8, Fisher Exact test (P<0.001) to determine bi-allelic single locus SNPs[21] and Chi-square test for bi-allelic segregation with 96% expected inbreeding. SNPs having two parents homozygous within and polymorphic between were extracted and missing loci were imputed with LB-impute [24] with parameter settings of -readerr 0.1 - genotypeerr 0.1 -window 7. Finally, SNP sites were removed if minor allele frequency (MAF) < 0.1, missing > 30% or heterozygosity > 6%. The TASSEL pipeline gave approximately 82% useable reads out of 2.15 billion reads. The overall alignment of 1,973,081 unique tags against the reference genome was 91% with a unique alignment of 37.8%. A total of 44,679 SNPs was discovered out of which 28,972 passed filtering. We then filtered RILs for missing data and heterozygosity resulting in 306 RILs and the two parents with suitable geneotypes. Finally, 8,797 SNPs were recovered after imputation with the additional filtering for MAF, missing and heterozygosity. All raw sequencing data for the Lakin x Fuller RIL population is available from NCBI SRA under accession number **SRP136362**.

Using traits directly from visual assessment and derived from image scoring by the K-net we calculated trait distributions, variance components, and best linear unbiased predictors in R statistical software [25] (Supplemental Information)

To model heading date from time series observations/predictions of percent heading, we fit a logistic growth curve model for each individual plot according to the function:

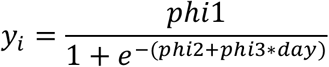

Where *y*_*i*_ is the is the *i*^*th*^ observation of heading percentage for a given plot. *phi1* is the asymptote maximum and was set to fixed at 100 for maximum percent heading. This model allows for different rates of development through heading as defined by *phi*2 and *phi*3. The independent variable *day* was calculated as the day of the year for observation *i*. The model was fit using nls function from nlme package [26]. To increase the robustness of fit we added points of 0 and 100 percent heading at 10, 20 and 30 days before the first and after the last visual measurements, respectively, corresponding to dates when all of the plots where not started and completely headed. The heading date for each plot was calculated as the date closest to 50% using the predict function in R at 0.1 day increments over the full range of the days.

For heading date, phi2 and phi3, we calculated broad-sense heritability on a line mean basis according to Holland *et al.*[27] for replicated trials of inbred lines (e.g. clonal species) in one location within one year as:

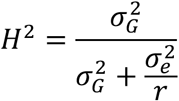

where 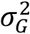 is the total genetic variance for entries in the trial, 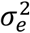 is the error variance and *r* is the number of replications. For heritability estimation across multiple years at one location as:

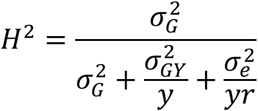

where 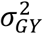 is the genotype by year variance and *y* is the number of years evaluated. Variance components were estimated by fitting mixed models in asreml package [28] in R. Models were fit with random effects for entry, year and replication within year and using a row-column autoregressive variance structure using the following model:

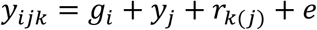

where, *y*_*ijk*_ is the observed plot-level phenotype, *g*_*i*_ is the random effect genotype effect of entry *i* distributed as iid where 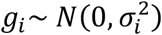, *y*_*j*_ is the random effect for year *j*, *r*_*k(j)*_ is the random effect of replication *k* within year *j*, and *e* is the residual variance partitioned with a two-dimensional autoregressive spatial structure (AR1 ⊗ AR1). Best linear unbiased estimates (BLUEs) were estimated for each entry within and across years by fitting the same model with entry as a fixed effect and using the predict function in R.

Following calculation of heading date, we observed a bimodal distribution of heading dates from the multi-year model BLUPs (**Supplemental Figure S4**). The distribution was delimited at 124 days to calculate the number of ‘early’ and ‘late’ lines. The number of lines in each group was fit to a *χ*^2^. test for two classes with probably of 0.75 and 0.25 according to a two-gene dominant epistasis model for inbred lines using the chisq.test function in R.

We tested for genetic association of heading date in the Lakin x Fuller population as measured by the logistic regression using a standard mixed model:

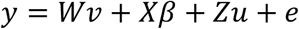

where *y* is the projected phenotype of heading date (50% intersect) or rate of heading (phi3) from the logisitic regression models. The use of a bi-parental population without population structure or kinship greatly simplified the equation to:

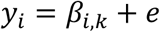

where *β*_*i,k*_ is the allele substitution effect for locus *k* in individual *i*; and *e* is residual error. Each marker effect was estimated using the lmer function in R and Bonferroni correction for multiple testing correction of experimental alpha of 0.05.

Following identification of significant marker association, we tested for two-way epistatic interactions for all markers that were associated with heading date using the model:

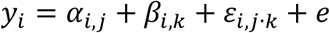

where, *y*_*i*_is the phenotype of the individual *i*, *α*_*i,j*_ is the allele substitution effect for locus *j* in individual *i*; *β*_*i,k*_ is the allele substitution effect for locus *k* in individual *i*; ɛ_*i,j·k*_ is the interaction between locus *j* and *k*; and *e* is residual error.

## ACKNOWLEDGEMENTS

We sincerely appreciate the assistance of Shuangye Wu in genotyping, Mark Lucas in data curation, and Haley Ahlers in graphics design, along with all members of the Wheat Genetics Lab at Kansas State University for project support, input and feedback. This work was supported by the National Science Foundation (NSF) Plant Genome Research Program (PGRP) (Grant No. IOS-1238187), the Kansas Wheat Commission and Kansas Wheat Alliance, the US Agency for International Development (USAID) Feed the Future Innovation Lab for Applied Wheat Genomics (Cooperative Agreement No. AID-OAA-A-13-00051), and by the NIFA International Wheat Yield Partnership (Grant No. 2017-67007-25933/project accession no. 1011391) from the USDA National Institute of Food and Agriculture. The opinions expressed herein are those of the author(s) and do not necessarily reflect the views of the U.S. Agency for International Development, the U.S. National Science Foundation, or the U.S. Department of Agriculture. The funders had no roll in study design, data collection or analysis.

## AUTHOR CONTRIBUTIONS

J.P. conceived and designed the study. B.E. managed the field trials and collected phenotypic data. X.W. developed the phenotyping platform and collected all image data. H.W. and R.P. analyzed images and developed neural networks. S.S. analyzed genetic data. J.P. directed the overall project, analyzed genetic and phenotypic data. J.P., X.W., H.W. and R.P. wrote the manuscript. All authors reviewed and approved the manuscript.

## COMPETING INTEREST

The authors declare no competing interests.

The plant materials tested in this study are public germplasm and/or were tested in accordance with local, national and international guidelines and legislation with the appropriate permissions and/or licenses for the study the present study.

