## Supplemental Document for "High-throughput phenotyping with deep learning gives insight into the genetic architecture of flowering time in wheat"

### SUPPLEMENTAL FIGURES

**Figure S1**. Example of geolocations of images from one camera (yellow points) marked on the field map (white polygons on the orthomosaic photo). GIS datasets (shape files) for all positioned images are included as supplemental data.

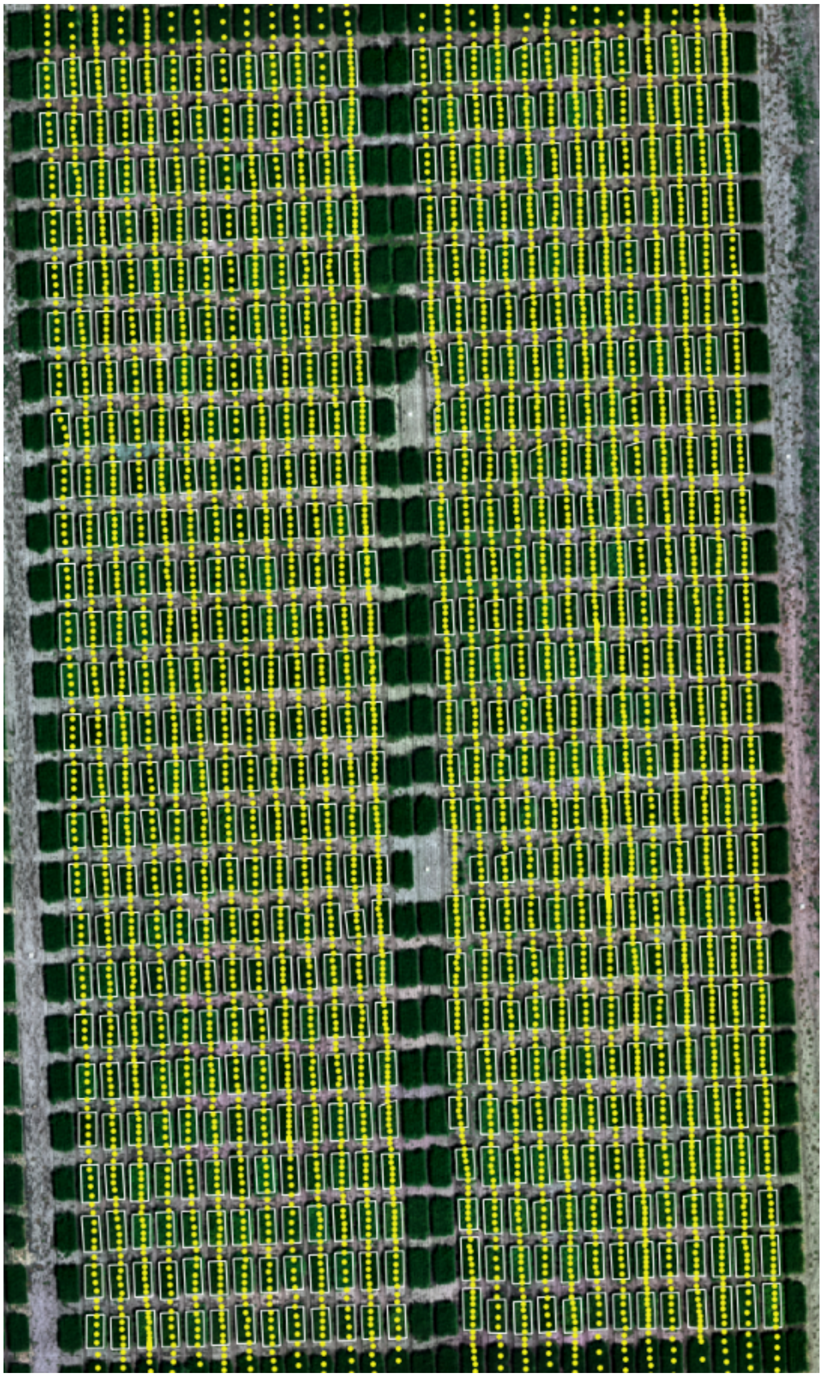

**Figure S2.** Example of (a) awnless and (b) awned phenotypes from the image dataset for the diversity panel.

a.

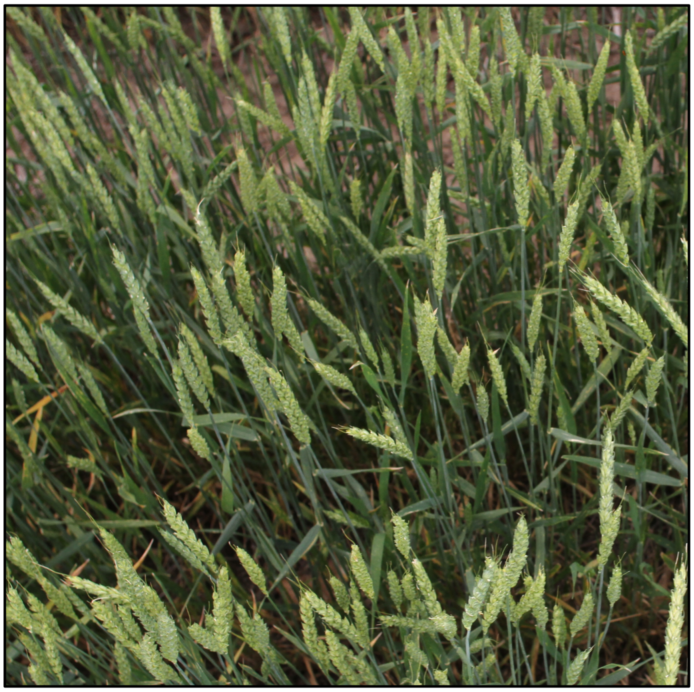

b.

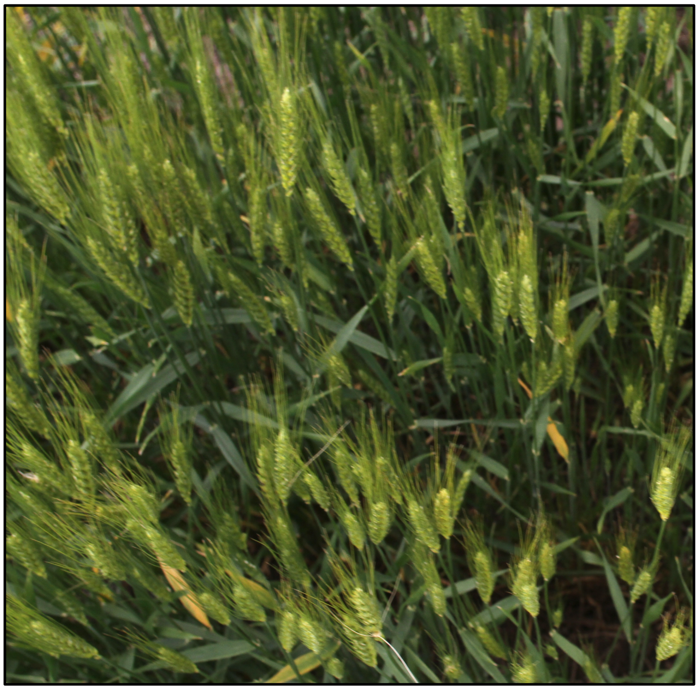

**Figure S3.** The confusion matrix for (a) training, (b) validation, and (c) testing the CNN for predicting percentage heading.

a.

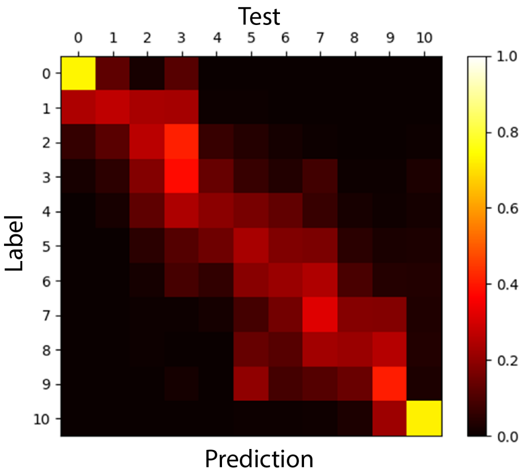


b.

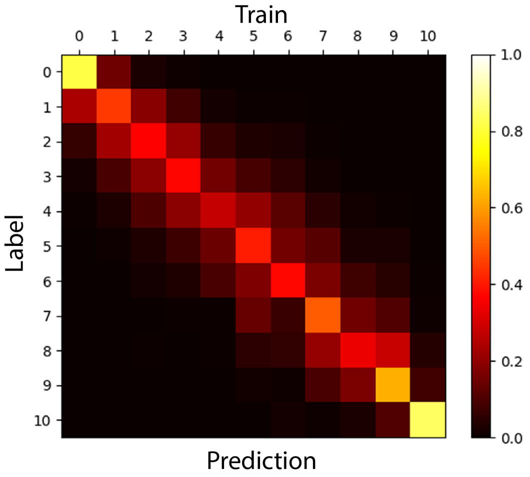


c.

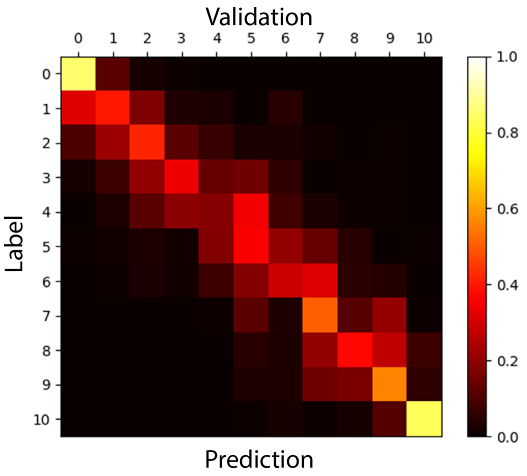

**Figure S4.** Phenotypic distribution of Lakin x Fuller recombinant inbred lines for heading date (day of year) in 2017 field trial.

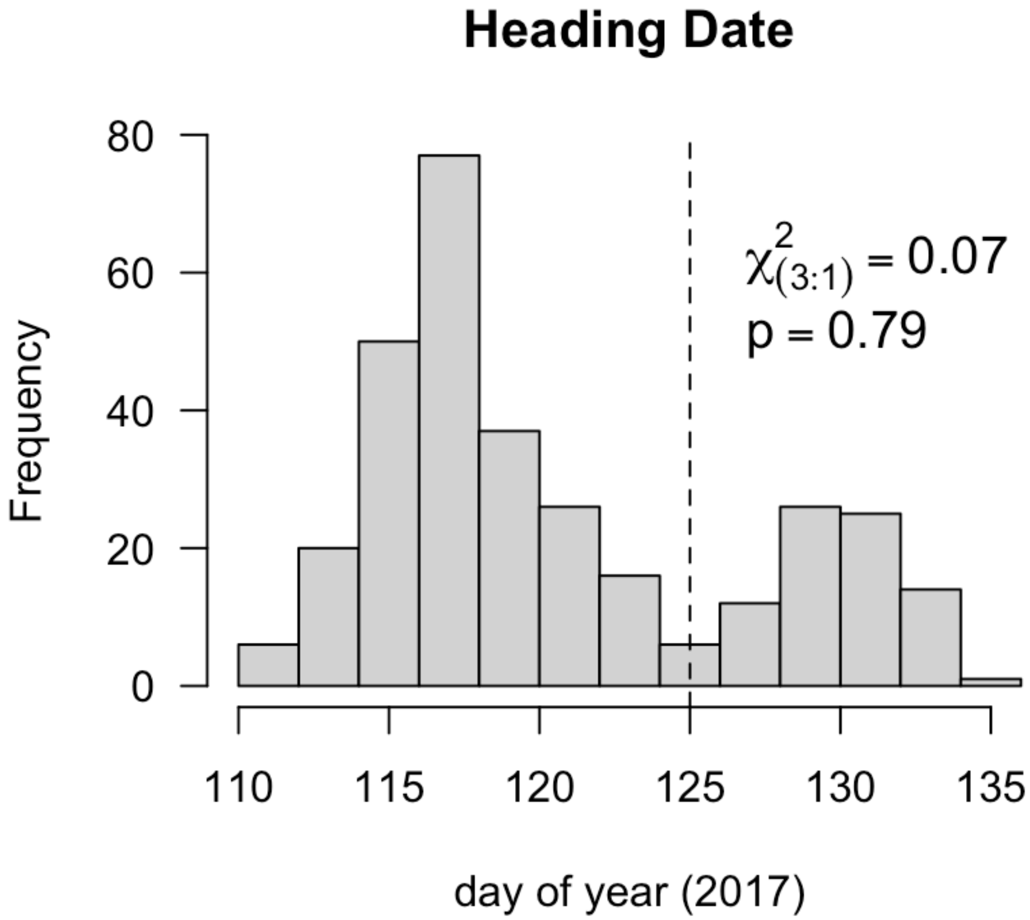

**Figure S5.** Genome wide association testing of phi2 (slope of logistic regression). No significant markers above Bonferroni multiple-test correction threshold.

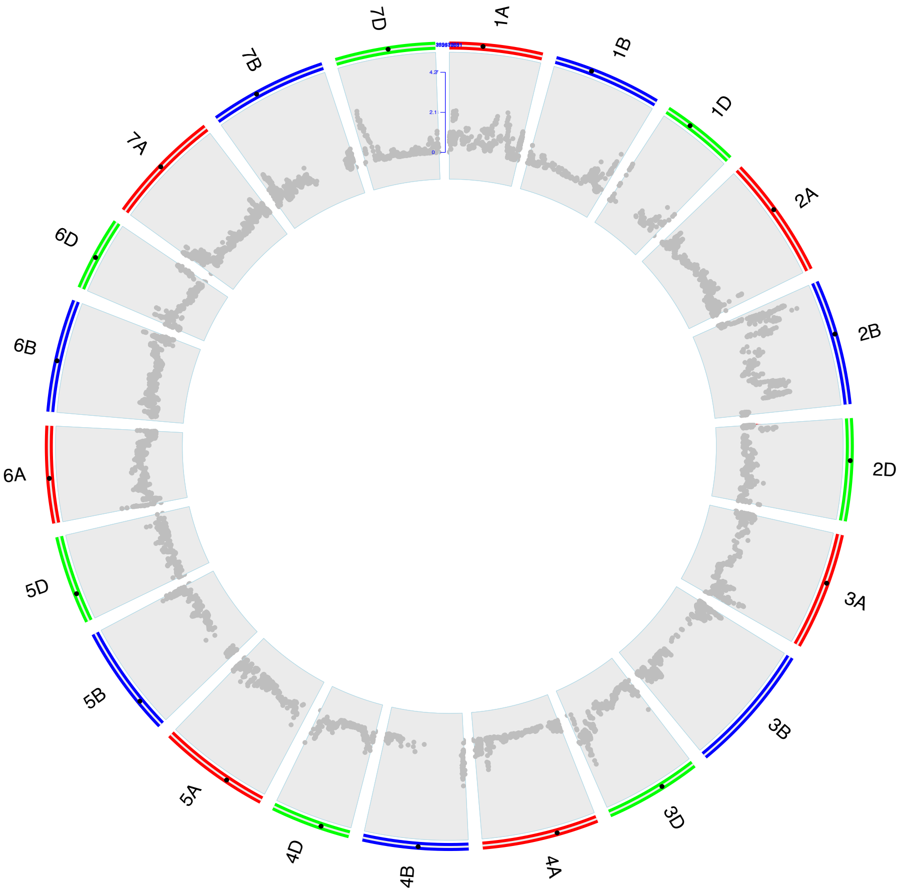

### SUPPLEMENTAL TABLES

**Table S1**. Individual image predictions (a) and accuracy (b) for awns phenotypes

a)

| | | Predicted | |
|---|---|---|---|
| | | AWNED | AWNLESS |
| **Visual** | AWNED | 10944 | 122 |
| | AWNLESS | 19 | 1419 |

b)

| | | Predicted | |
|---|---|---|---|
| | | AWNED | AWNLESS |
| **Visual** | AWNED | 0.988 | 0.011 |
| | AWNLESS | 0.013 | 0.986 |

**Table S2**. Supplemental dataset table of field plot entries and field design for Diversity Panel and Lakin x Fuller Recombinant Inbred Line (RIL) populations in 2016 and 2017.

**Supplemental Table S3**. Full details of each layer in the WheatNet.

| Net | Block | Layer | function | in | out | parameters | Layer | function | in | out | parameters |
|---|---|---|---|---|---|---|---|---|---|---|---|
| ResNet | | layer1 | conv_7x7 | 9xNx3x224x224 | 9xNx64x112x112 | 9408 | | | | | |
| | | | bn_2D | 9xNx64x112x112 | 9xNx64x112x112 | 128 | | | | | |
| | | | relu | 9xNx64x112x112 | 9xNx64x112x112 | | | | | | |
| | | | maxpool_3x3 | 9xNx64x112x112 | 9xNx64x56x56 | | | | | | |
| | Block_1 | layer2 | conv_3x3 | 9xNx64x56x56 | 9xNx64x56x56 | 73728 | downsampler | conv_1x1 | 9xNx64x56x56 | 9xNx64x56x56 | 4096 |
| | | | bn_2D | 9xNx64x56x56 | 9xNx64x56x56 | 128 | | | | | |
| | | | relu | 9xNx64x56x56 | 9xNx64x56x56 | | | | | | |
| | | | drop_out | | | | | bn_2D | 9xNx64x56x56 | 9xNx64x56x56 | 128 |
| | | layer3 | conv_3x3 | 9xNx64x56x56 | 9xNx64x56x56 | 36864 | | | | | |
| | | | bn_2D | 9xNx64x56x56 | 9xNx64x56x56 | 128 | | | | | |
| | | | drop_out | | | | | | | | |
| | Block_2 | layer4 | conv_3x3 | 9xNx64x56x56 | 9xNx128x28x28 | 73728 | downsampler | conv_1x1 | 9xNx64x56x56 | 9xNx128x28x28 | 8192 |
| | | | bn_2D | 9xNx128x28x28 | 9xNx128x28x28 | 256 | | | | | |
| | | | relu | 9xNx128x28x28 | 9xNx128x28x28 | | | | | | |
| | | | drop_out | | | | | bn_2D | 9xNx128x28x28 | 9xNx128x28x28 | 256 |
| | | layer5 | conv_3x3 | 9xNx128x28x28 | 9xNx128x28x28 | 147456 | | | | | |
| | | | bn_2D | 9xNx128x28x28 | 9xNx128x28x28 | 256 | | | | | |
| | | | drop_out | | | | | | | | |
| | Block_3 | layer6 | conv_3x3 | 9xNx128x28x28 | 9xNx256x14x14 | 294912 | downsampler | conv_1x1 | 9xNx128x28x28 | 9xNx256x14x14 | 32768 |
| | | | bn_2D | 9xNx256x14x14 | 9xNx256x14x14 | 512 | | | | | |
| | | | relu | 9xNx256x14x14 | 9xNx256x14x14 | | | | | | |
| | | | drop_out | | | | | bn_2D | 9xNx256x14x14 | 9xNx256x14x14 | 512 |
| | | layer7 | conv_3x3 | 9xNx256x14x14 | 9xNx256x14x14 | 589824 | | | | | |
| | | | bn_2D | 9xNx256x14x14 | 9xNx256x14x14 | 512 | | | | | |
| Summarization | | layer8 | avgpooling_7x7 | 9xNx256x14x14 | 9xNx256x1x1 | | | | | | |
| | | layer9 | fc | Nx9x256 | Nx9x11 | 2816 | | | | | |
| | | layer10 | avgpooling_9x1 | Nx9x11 | Nx1x11 | | | | | | |
| | | layer11 | fc | Nx11 | Nx11 | 121 | | | | | |
| | | | | | | 1276729 | | | | | |

**Table S4.** Analysis of variance for full interaction model of Ppd-D1, Ppd-B1 and locus on 1B.

test = lm(Y~g.2B*g.2D*g.1B)

Analysis of Variance

| | Df | Sum Sq | Mean Sq | F value | Pr(>F) |
|---|---|---|---|---|---|
| **g.2B** | 1 | 2486 | 2486 | 259.7034 | < 2.2e-16 *** |
| **g.2D** | 1 | 5587.1 | 5587.1 | 583.6726 | < 2.2e-16 *** |
| **g.1B** | 1 | 241.7 | 241.7 | 25.2465 | 8.887e-07 *** |
| **g.2B:g.2D** | 1 | 1026.8 | 1026.8 | 107.2628 | < 2.2e-16 *** |
| **g.2B:g.1B** | 1 | 1.5 | 1.5 | 0.155 | 0.694053 |
| **g.2D:g.1B** | 1 | 99.3 | 99.3 | 10.3704 | 0.001428 ** |
| **g.2B:g.2D:g.1B** | 1 | 48.7 | 48.7 | 5.0921 | 0.024787 * |
| **Residuals** | 287 | 2747.3 | 9.6 | | |

Signif. codes:  0 ‘***’ 0.001 ‘**’ 0.01 ‘*’ 0.05 ‘.’ 0.1 ‘ ’ 1
